# Regulation of touch dependant *de novo* root regeneration in Arabidopsis

**DOI:** 10.1101/2020.11.04.367714

**Authors:** Anju Pallipurath Shanmukhan, Mabel Maria Mathew, Mohammed Aiyaz, Abdul Kareem, Dhanya Radhakrishnan, Kalika Prasad

## Abstract

The versatile regeneration capability of leaves enable even a detached Arabidopsis leaf to yield two kinds of regenerative responses namely, wound healing at the cut end in form of callus formation or de novo root regeneration (DNRR). Using various experimental approaches, we show that the factor favouring DNRR over callus formation seems to be a mechanical cue, possibly touch, at the cut end of the detached leaf. Here, we show that the forced expression of a PLETHORA transcription factor can bypass the need for touch to initiate DNRR. Our findings provide a genetic frame-work for touch dependant DNRR and suggest that a core PLT transcription regulatory module acts in response to mechano-sensing stimuli.

---

Among several of the plant’s lateral organs, leaves show highly versatile yet efficient regenerative responses. Regeneration in leaves may be natural, mechanical injury-induced, or tissue culture-mediated. In natural regeneration, an entire plant can regenerate from the leaf without hormonal supplements, for example various Kalanchoe species(Smith, Figueiredo and Van Wyk, 2019). In tissue culture-mediated regeneration, small leaf explants can give rise to entire shoot and/root system via an intermediary callus stage, but in the presence of hormonal supplements. Apart from these, the incised mid-vein of an undetached growing leaf and the cut end of a detached leaf exhibit regenerative responses, both of which fall under mechanical injury-induced regeneration. Although the mid-vein regeneration in growing leaves was investigated only recently, mechanical injury-induced regenerative responses at cut end of detached leaf has been studied for several years now(Chen *et al.*, 2014; Bustillo-Avendaño *et al.*, 2018; Xu, 2018; Radhakrishnan *et al.*, 2020). Adventitious roots arise from the cut site of the detached leaf, be it the base of leaf blade or the petiole via the process of *de novo* root regeneration (DNRR)(Chen *et al.*, 2014; Bustillo-Avendaño *et al.*, 2018). This ability of part of a tissue to give rise to an organ, whose identity is different from its parent tissue is rather intriguing. However, DNRR is not the only response observed at the cut end; wound healing in the form of callus formation is seen at the cut end of leaves that do not undergo DNRR. With the previously available data, it is unclear if the decision to make callus or DNRR is random or if any external inductive cues favour one phenomenon over the other. It is therefore imperative that we investigate this differential regenerative response to the same injury in the same organ.

We repeated the previously established DNRR assay using whole leaves with petioles (Bustillo-Avendaño *et al.*, 2018). When the leaves were placed abaxial-side down with the cut end of petiole touching the surface of the hormone free-solid Murashige and Skoog (MS)-Agar medium (referred to as MS-Agar media hereafter), we observed DNRR which is consistent with the previous studies (Fig. 1, A,A’,B and B’)(Bustillo-Avendaño *et al.*, 2018)). In our hand, the success rate of DNRR was only 30% and the remaining 70% of the leaves showed neither DNRR nor callus formation(n=62) (Fig 1B’). However, when the leaves were placed adaxial-side down with the petiole in the air and not touching the surface of media, DNRR failed to occur in all the samples, but callus formation was observed at the cut end in 62% (n=50) of the samples (Fig. 1, C,C’,D and D’). At a glance, three factors appear to be different between the two cases: (i) nutrient availability at cut end (minimal MS and sucrose), (ii) the orientation of the leaf on the MS-Agar media (adaxial/abaxial), and (iii) physical touch of the cut end to the solid agar surface. Upon closer examination, the factor that separates the two explants and presumably leading to the distinct regenerative responses seems to be, touch. The explants that produced DNRR had its cut end touching the surface of the MS-Agar media while those explants that had only callus formation were devoid of touch. To prevent absorption of nutrients by the cut end from interfering with this observation, we first carried out a simple split plate experiment where the top half of the plate contains MS-Agar media, while the bottom half contains nutrient-free solid agar media (referred to as agar-only media hereafter). The leaves were placed abaxial side down, such that the top half of the leaf touches the media with nutrients and the bottom half with the petiole touches the nutrient-free media (Fig. 1E). Since detached leaves on agar-only media showed yellowing and withered away (Fig. S1A), we allowed the top half of leaf to touch the MS-agar media to allow minimal nutrient transport for its sustenance and growth. Interestingly, 20.78% (n=154) of leaf explants produced DNRR from the cut end that touched the agar-only media (Fig. 1F), while the remaining leaf explants produced neither DNRR nor callus formation. Secondly, to examine if the orientation of the leaf on the media plays a role, we placed the leaf adaxial side down on an MS-Agar media while gently pressing down the leaf ensuring that the cut end touches the media (Fig. 1G). We noticed that 34.2 % (n=38) of leaf explants exhibited DNRR (Fig. 1H). Thirdly, we designed an experiment combining the first and second experiments, where the detached leaf was placed adaxial-side down on the half MS-Agar media, with its petiole being sandwiched between a parafilm strip and a block of agar-only media(Fig. 1I). The experimental set up ensures that the leaf is oriented adaxial-side down, devoid of nutrients at cut end and that the cut end is in touch with a nutrient-free agar block. We noticed that 17.86% (n=56) of leaf explants exhibited DNRR (Fig. 1J). Collectively, the results were in agreement with our hypothesis, that touch is the major factor distinguishing the two regenerative responses at the cut end of a detached leaf namely, DNNR from callus formation.

**Figure 1:**
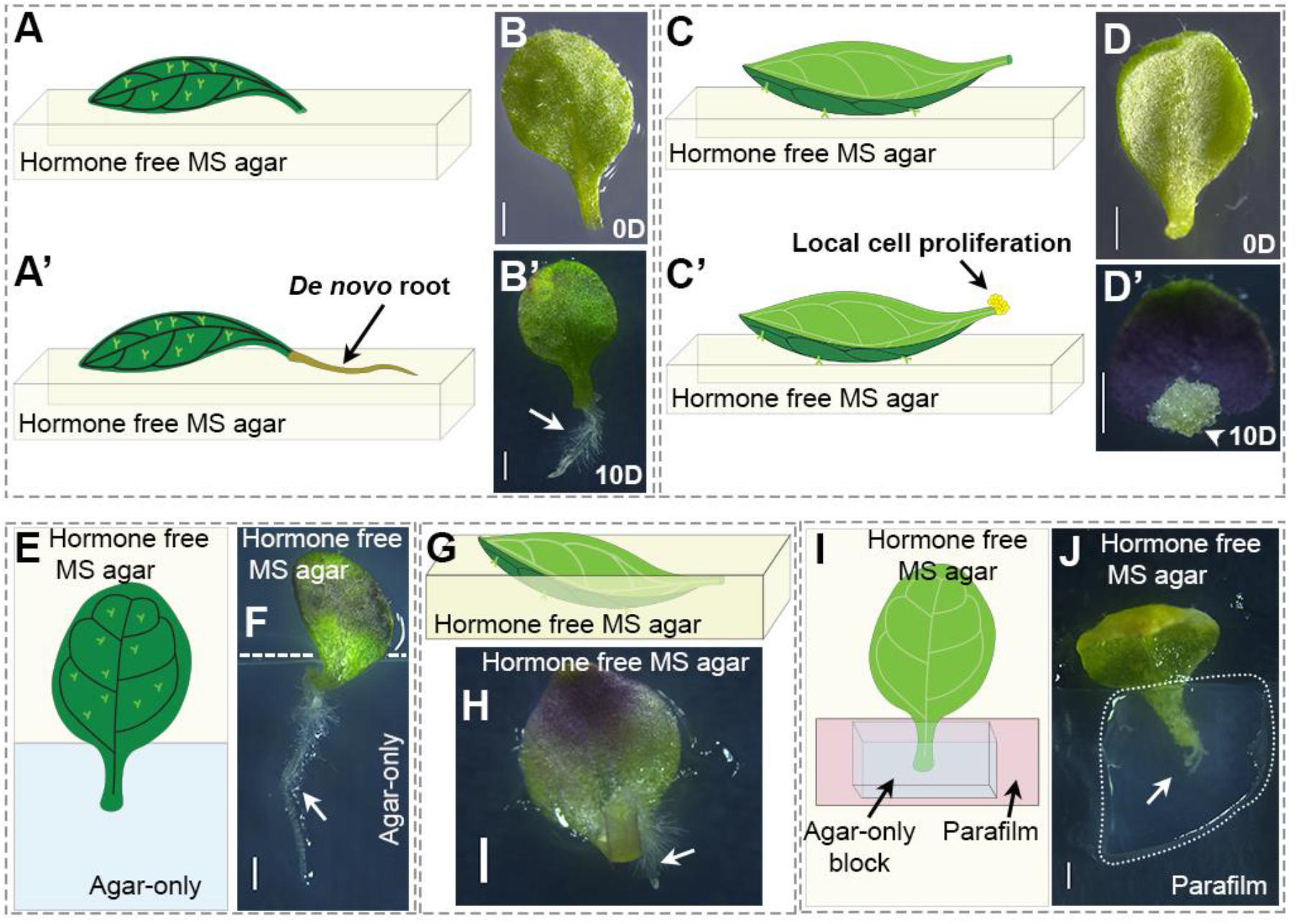
Wound healing response and touch-dependant de novo regeneration at the cut end of a detached leaf (A-I): (A,A’) A detached leaf when placed abaxial side down on the hormone-free solid MS-Agar media (referred as MS-Agar media hereafter) results in the formation of *de novo* root. (B,B’)The stereo-microscopic images of the detached leaf placed abaxial side down that regenerated *de novo* roots. (C,C’)A detached leaf when placed adaxial side down on the MS-Agar media results in the formation of *de novo* root. (D,D’)The stereo-micrographs of the detached leaf placed adaxial side down that regenerated *de novo* roots. (E) Schematic depicting a “split-plate” where top half of MS-Agar medium is insulated from hormone-free solid agar-only medium (referred as Agar-only medium hereafter). The leaf is placed abaxial side down with its distal end touching the MS-Agar medium and its cut end touching the Agar-only medium. (F) Stereo-micrographs of the leaf showing DNRR on the split plate. (G) Schematic showing the detached leaf being pressed into the media with its adaxial side down. (H) Stereo-micrograph of the leaf showing DNRR after being pressed into the medium. (I) Schematic illustrating the experimental set up where the cut end touches Agar-only media but insulated from MS-Agar media. Here, the detached leaf is placed abaxial side down on MS-Agar medium, and the cut end is sandwiched between a thin parafilm strip and an Agar-only block. (J) Stereo micrograph of leaf showing DNRR after the cut end being sandwiched between parafilm and Agar-only block. The black and while arrows indicate *de novo* regenerated root. Scale bars represent 1mm.

We then explored the genetics underlying this touch-dependant DNRR. Due to their indispensible and established role in tissue culture and injury-induced regeneration in Arabidopsis, *PLETHORA 3 (PLT3), PLT5, and PLT7* were our first choice for investigation (Fig. 2, A-H and Fig. S1, B-O). Upon examining the expression pattern of PLT7 using WT/*PLT7::PLT7-YFP* we found that the several cells near the cut expressed PLT7-YFP, at a low intensity by 24 hour when the cut end continuously touched the MS-Agar media; by 48hours the YFP expression became more prominent (Fig. 2, A-D). However, when the cut end failed to touch the MS-Agar media, only few cells at the cut end expressed PLT7-YFP and at very low intensity by 24 hours, and by 48hours the YFP expression still remained faint and confined to very few cells (Fig. 2, E-H). It should be noted that all the leaf explants were cultured on hormone-free solid MS-agar media hereafter. Leaf explants from *plt3,5,7* mutant failed to yield DNRR despite the cut end of the petiole being in contact with the solid MS-Agar media, while over-expression(OE) of PLT7 (WT/*35S::PLT7-GR*) in Wild type background induced excessive DNRR (Fig. 2, I and J, Fig. S1, P and Q). Surprisingly, PLT7-OE also induced DNRR at a frequency of 52.2% (n=46) even when the cut end does not touch the media (Fig. 2, K and L). Thus, PLT7-OE seems to over-ride the need for touch suggesting that PLT7 is necessary and sufficient to induce DNRR. Until now, PLT3,5,7 has been known to act through two different transcriptional regulatory modules during regeneration: (i) a two-step mechanism in which PLT3,5,7 independently activates PLT1,2 and CUC2 sequentially for tissue culture-induced *de novo* shoot regeneration, and (ii) a PLT-CUC2 regulatory axis which acts in a coherent feed-foreword loop to upregulate local auxin biosynthesis gene YUCCA4 during mechanical injury-induced vein regeneration in an undetached growing leaf (Kareem *et al.*, 2015; Radhakrishnan *et al.*, 2020). We probed, if either of the two regulatory modules could plausibly act during DNRR at the cut end. Live imaging using fluorescent reporter lines showed that PLT1 and PLT2 upregulated their expression during DNRR (Fig. S2A,A’,C and C’). Additionally, expression of CUC2 was also observed, albeit rarely and at low level (Fig. S2, E and E’).

**Figure 2:**
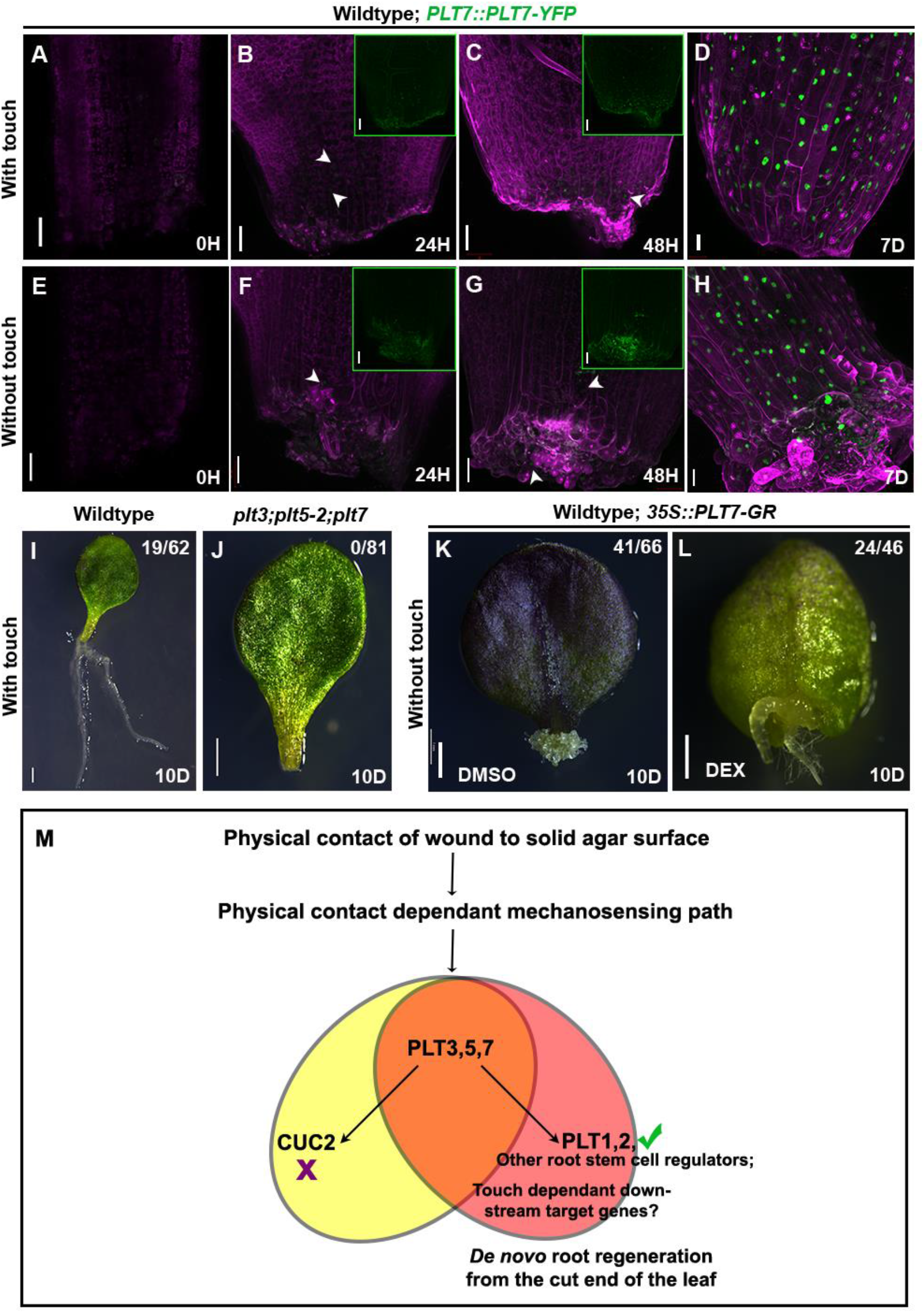
PLT3,5,7 are necessary and sufficient for touch mediated *de novo* root regeneration: (A-H) *PLT7::PLT7-YFP* expression (green) expression marked by white arrow heads when the cut end is in continuous contact with the MS-Agar medium (A-D) and when the cut end fails to touch the medium (E-H). Insets in B,C,F,G shows the YFP-channel. (I,J) Wild type (WT) leaf explants exhibit DNRR(I) while *plt3;plt5-2;plt7* mutant(J) shows neither callus formation nor DNRR even when the cut end touches the MS-Agar medium. (K, L) Over expression with *35S::PLT7-GR* yields DNRR even when the cut end fails to touches the MS-Agar medium. (M) Flow-chart illustrating the regulatory module that influence touch dependant DNRR from cut end of a detached leaf. Scale bars:50μm (A-H), 1mm (I-L). Magenta colour denotes chlorophyll autofluoresence and propidium iodide. H: hours post cut, D: days post cut. Brightness and contrast has been adjusted for B,C,F and G insets.

Unlike in Wildtype (WT), PLT1, PLT2 as well as CUC2 remain undetectable in the cut ends of *plt3,5,7* mutant leaf explants (Fig. S2, B,B’,D,D’,F,F’). *plt3,5,7* mutants are defective in lateral root (LR) outgrowth. However, reconstitution of PLT1 in *plt3,5,7* mutant (*plt3,5,7/PLT7::PLT1-YFP*) rescued the root regeneration when petiole touches the medium, whilereconstitution of CUC2 in *plt3,5,7* mutant (*plt3,5,7/PLT5::CUC2-YFP*) failed to do so (Fig. S3). Thus, PLT3,5,7-CUC2 regulatory module seems to be dispensable while, PLT3,5,7 mediated activation of PLT1 appears to control DNRR in response to touch at the cut end. Very few of the *plt3,5,7* mutant leaf explants showed DNRR upon PLT1 reconstitution as opposed to the DNRR in 30% WT explants suggesting a partial recovery (Fig S3). Here, the low DNRR frequency is in contrast to the complete rescue of LR out-growth in *plt3,5,7* mutant samples reconstituted with PLT1(Du and Scheres, 2017). Moreover, reconstitution of only PLT1 and not other root stem cell regulators such as WOX5 could trigger DNRR (Fig S3). Taken together, our results suggest that PLT mediated touch-dependant DNRR follows a mechanism distinct from that of PLT mediated regulation of LR outgrowth. PLT3,5,7 regulated DNRR likely requires additional root stem cell regulators, and this notion is in line with the previous reports that the cumulative loss of function of PLT1, PLT2 and SHR severely impair DNRR(Bustillo-Avendaño *et al.*, 2018). It is plausible that, in addition to root stem cell regulators there are additional downstream targets which are activated by PLT3,5,7 in response to mechano-sensitive cues (Fig. 2M).

Recently, it was shown that regeneration of specific cell types in root was influenced by osmotic pressure, suggesting that mechanical signalling pathways can be instrumental in regeneration(Hoermayer *et al.*, 2020). Therefore, it is possible that mechanical signals can instruct DNRR. Mechanical signal transduction can be triggered when the mechano-receptors on cells perceive stimuli such as gravity, wind, turgor pressure and in this case, touch. At present, it is likely that the touch induced mechano-signal transduction activate the regulatory module for fate switch from leaf to root rather indirectly. Here, we show that touch-dependant DNRR from detached leaf acts atleast in part through the root stem cell regulators PLT3,5,7 and PLT1(Fig. 2M). It will be interesting to unravel how mechano-sensing impacts a PLT-regulated genetic frame-work in the event of DNRR from a detached leaf.

## Acknowledgements & Funding

K.P. acknowledges Department of Biotechnology (DBT), Government of India [grant BT/PR12394/AGIII/103/891/2014] and Department of Science and Technology, Science and Engineering Research Board (DST-SERB), Government of India [grant EMR/2017/002503/PS] for funding and also acknowledges the Indian Institute of Science Education and Research Thiruvananthapuram (IISER-TVM) for infrastructure facilities.

A.P.S. is a recipient of Council of Scientific and Industrial Research (CSIR) fellowship. D.R. and M.M.M acknowledge University Grants Commission (UGC) fellowship. A.K. was recipient of Indian Institute of Science Education and Research-Thiruvananthapuram fellowship. M.A. acknowledges Department of Biotechnology (DBT), Ministry of Science and Technology, Government of India for the DBT-Post Doctoral Fellowship (DBT-RA Program).

