## Supplementary 1 for "Regulation of touch dependant *de novo* root regeneration in Arabidopsis"

### Supplementary figures and legends

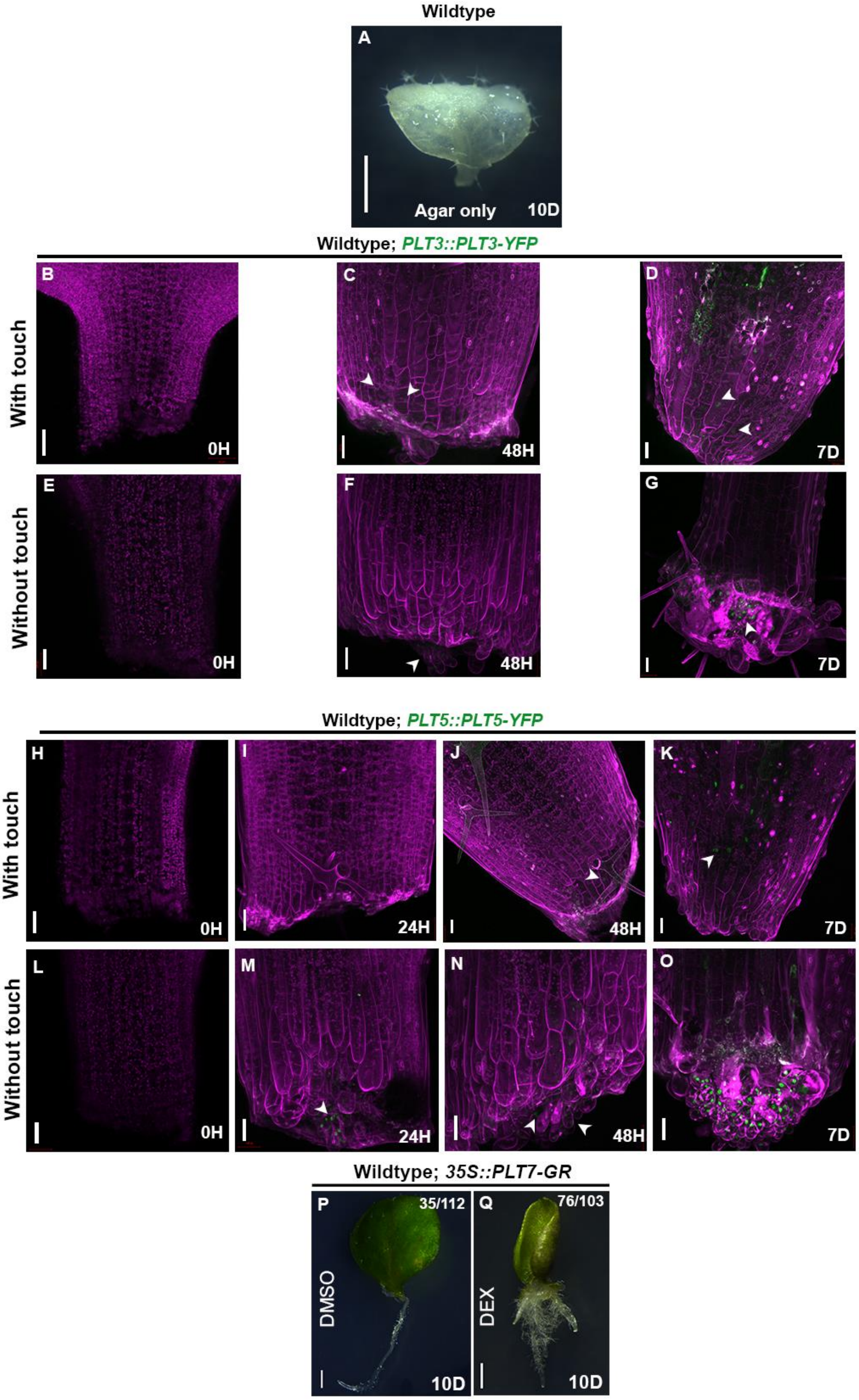
Wildtype
A
Agar only
10D
Wildtype; PLT3::PLT3-YFP
With touch
Without touch
B
0H
C
48H
D
7D
E
0H
F
48H
G
7D
Wildtype; PLT5::PLT5-YFP
With touch
Without touch
H
0H
I
24H
J
48H
K
7D
L
0H
M
24H
N
48H
O
7D
Wildtype; 35S::PLT7-GR
P
35/112
DMSO
10D
Q
76/103
DEX
10D

**Supplementary figure 1: (**A) Response of detached leaf when placed on agar-only media. (B-D) *PLT3::PLT3-YFP*(green) expression(marked by white arrowhead) at cut end of leaf when it is in contact with media. (E-G) *PLT3::PLT3-YFP* (green) expression at cut end of petiole when it is not in contact with media. (H-K) *PLT5::PLT5-YFP* expression at cut end of petiole when it is in contact with media. (L-O) Expression of *PLT5::PLT5-YFP* in cut ends of detached leaf when there is no contact with media. (P,Q) Overexpression of *35S::PLT7-GR* causes multiple root formation from the cut end. Scale bar: 50 µm (B-O), 1mm (A,P,Q). Magenta colour denotes chlorophyll autofluoresence and propidium iodide. H: hours post cut, D: days post cut.

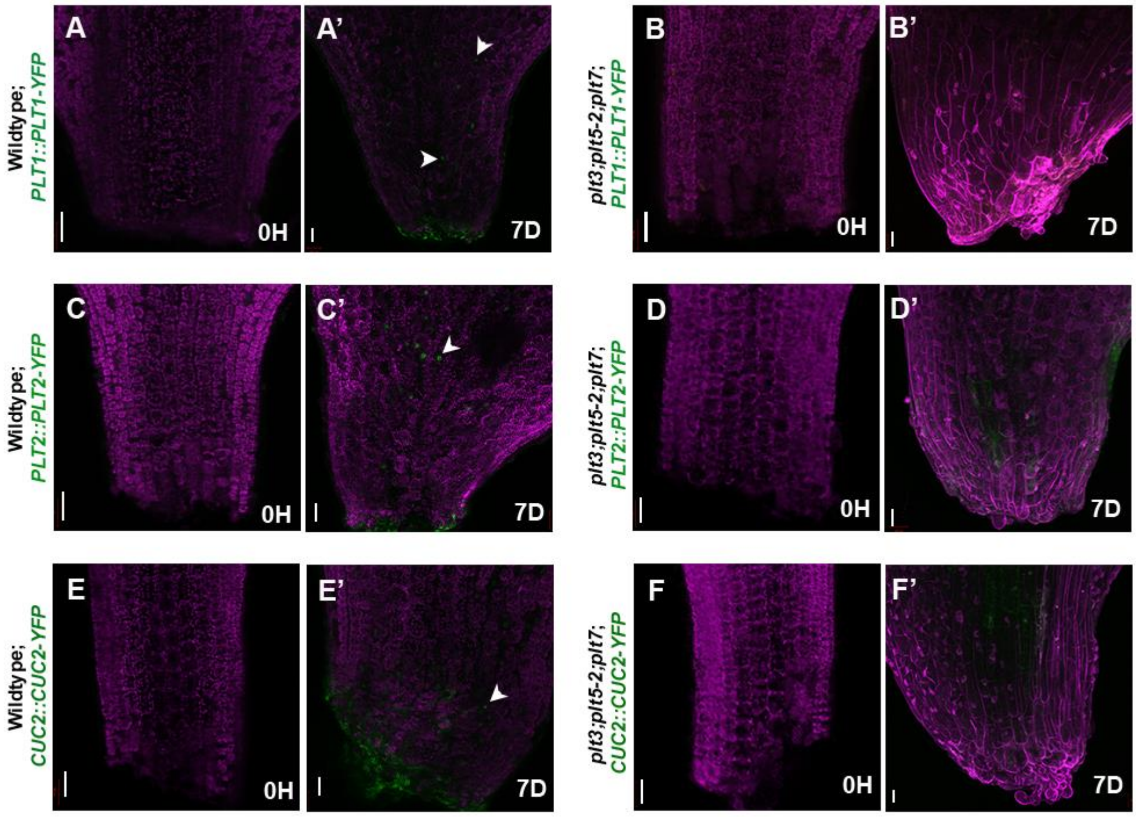


**Supplementary figure 2: (**A,A', B,B') Expression of *PLT1::PLT1-YFP* (green) in detached leaves of Wildtype (A,A') and *plt3,plt5-2,plt7* (B,B') mutant as marked by white arrowhead. (C,C', D,D') Expression pattern(arrowhead) of *PLT2::PLT2-YFP* (green) in detached leaves of Wildtype (C,C') and *plt3,plt5-2,plt7* mutant(D,D'). (E,E', F,F') *CUC2::CUC2-YFP* expression (green) in detached leaves of Wildtype (E,E') and *plt3,plt5-2,plt7* mutant (F,F'). Scale bar: 50µm, Magenta colour denotes chlorophyll autofluorescence and propidium iodide. H: hours post cut, D: days post cut.

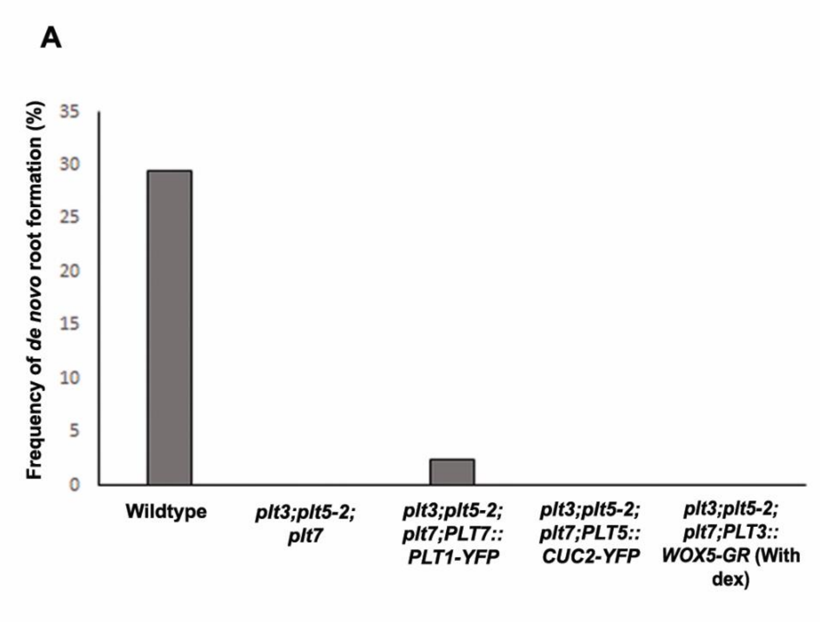


**Supplementary figure 3:** A) Graph showing comparison of frequency of *de novo* root regeneration in Wildtype, *plt3,plt5-2,plt7, plt3,plt5-2,plt7 PLT7::PLT1-YFP, plt3,plt5-2,plt7 PLT5::CUC2-YFP, plt3,plt5-2,plt7 PLT3::WOX5-GR* (with dex).

### Materials and methods

Growth conditions

Arabidopsis thaliana seeds were sterilised using 70% ethanol and 20% bleach, followed by seven washes with sterile distilled water. Seeds were plated on half-strength Murashige-Skoog (MS) medium (pH 5.7) with 0.7% agar and grown vertically under 45 µmol/m2/s continuous white light at 22°C and 70% relative humidity.

Regeneration assays

All the plants were grown on hormone-free half-strength MS-Agar medium (Sigma-Aldrich). To study de novo regeneration responses in detached leaves, 7 day old seedlings were chosen and the first pair of leaves were excised using Vannas straight scissors and placed on hormone free half strength MS agar media (with 0.7% agar). The detached leaves with petiole were placed either the abaxial side or adaxial side facing the media. The leaves where abaxial side faces media/cut end is in contact with media produced de novo roots. When the cut end of leaf is protruding out of media/adaxial side facing media, local wound healing response in the form of callus was observed. Both the regeneration responses were scored 10 days after excision.

*De novo* root regeneration assay

For *de novo* root formation, the leaf explants with petiole were placed on half strength MS-Agar medium (containing 0.7% agar) with the cut end of the petiole touching the medium (abaxial side of the leaf facing media).

1. Split plate experiment

Hormone free solid MS-Agar (0.7%) was poured into top half of a pre-sterilized square petri-plate (HIMEDIA) that was split down the middle using a pre-sterilized insulator. The bottom half of the plate was filled with hormone-free solid Agar-only media (0.7%). The detached leaves were placed abaxial side down across the middle split such that, the distal region of the leaf was in contact with the MS-Agar media and the cut end of the leaf with its petiole touched the Agar-only media. The leaf explants were cultured in the plate and scored for regeneration on the 10$^{th}$ day.

2. Leaf pressed into media

Hormone free solid MS-Agar (0.7%) was poured into a pre-sterilized square petri-plate (HIMEDIA) and allowed to cool. The detached leaves were placed adaxial side down and was pressed down into the media such that the cut end of the leaf touched the media. The leaf explants were cultured in the plate and scored for regeneration on the 10$^{th}$ day.

3. Cut end of leaf sandwiched between parafilm and Agar block

Hormone free solid MS-Agar (0.7%) was poured into a pre-sterilized square petri-plate (HIMEDIA) and allowed to cool and a thin strip of parafilm strip was placed across the MS-Agar media. The detached leaves were placed adaxial side down and ensuring that only the distal region of the leaves were in contact with the MS-Agar media and the cut end and petiole were insulated from it by the parafilm strip. Hormone-free Agar-only blocks were placed on the cut end of the leaves, thereby sandwiching the cut ends between the parafilm and the Agar-only block. The leaf explants were cultured in the plate and scored for regeneration on the 10$^{th}$ day.

Plant materials

Arabidopsis thaliana ecotype Columbia (Col-0) was used as wild type for the study. The genetic backgrounds and translational fusion constructs used for the study were *plt3;plt5-2,plt7* and *PLT1::PLT1-vYFP, PLT2::PLT2-vYFP, PLT3::PLT3-vYFP, PLT5::PLT5-vYFP, PLT7::PLT7-vYFP, CUC2::CUC2-vYFP and 35S::PLT7-GR* as described in Radhakrishnan et al., 2020 and *PLT7::PLT1-vYFP* as described in Kareem et al., 2015, *PLT5::CUC2-vYFP* was generated using Multigateway recombination system.

Microscopic imaging

Brightfield and confocal laser-scanning microscopy imaging were performed as described previously in Kareem et al., 2015. Confocal imaging of leaf samples were performed using Zeiss LSM 880 confocal laser-scanning microscope and brightfield images were acquired using Leica M205 FA fluorescence stereo microscope. The cell boundaries of detached leaf samples were stained using 10 μg/ml propidium iodide (Sigma-Aldrich) for confocal imaging. Images were acquired using 10× air, 20× air objectives. The images were processed using Zeiss ZEN black software. Schematics were drawn using Adobe illustrator CC 2018. Images were compiled using Adobe Photoshop CS6.
